# GTDrift: A resource for exploring the interplay between genetic drift, genomic and transcriptomic characteristics in eukaryotes

**DOI:** 10.1101/2024.01.23.576799

**Authors:** Florian Bénitière, Laurent Duret, Anamaria Necsulea

## Abstract

We present GTDrift, a comprehensive data resource that enables explorations of genomic and transcriptomic characteristics alongside proxies of the intensity of genetic drift in individual species. This resource encompasses data for 1,506 eukaryotic species, including 1,413 animals and 93 green plants, and is organized in three components. The first two components contain approximations of the effective population size, which serve as indicators of the extent of random genetic drift within each species. In the first component, we meticulously investigated public databases to assemble data on life history traits such as longevity, adult body length and body mass for a set of 979 species. The second component includes estimations of the ratio between the rate of non-synonymous substitutions and the rate of synonymous substitutions (*dN/dS*) in protein-coding sequences for 1,324 species. This ratio provides an estimate of the efficiency of natural selection in purging deleterious substitutions. Additionally, we present polymorphism-derived *N*_e_ estimates for 66 species. The third component encompasses various genomic and transcriptomic characteristics. With this component, we aim to facilitate comparative transcriptomics analyses across species, by providing easy-to-use processed data for more than 16,000 RNA-seq samples across 491 species. These data include intron-centered alternative splicing frequencies, gene expression levels and sequencing depth statistics for each species, obtained with a homogeneous analysis protocol. To enable cross-species comparisons, we provide orthology predictions for conserved single-copy genes based on BUSCO gene sets. To illustrate the possible uses of this database, we identify the most frequently used introns for each gene and we assess how the sequencing depth available for each species affects our power to identify major and minor splice variants.

## Introduction

Genetic drift refers to stochastic fluctuations in allele frequencies within a population across successive generations. These fluctuations arise due to the inherently random sampling of individuals that reproduce and pass on their alleles to subsequent generations (Wright, 1929; Graur and Li, 2000). Population genetics principles state that the ability of natural selection to promote beneficial mutations or eliminate deleterious mutations depends on the intensity of selection (*s*) relative to the power of genetic drift (defined by the effective population size, *N* _e_): if the selection coefficient is sufficiently weak relative to drift (|*N* _e_*s*| *<* 1), alleles behave as if they are effectively neutral (Kimura *et al*., 1963; Ohta, 1973). Thus, random drift sets an upper limit on the efficiency of selection. This limit is called the “drift barrier” (Lynch, 2007, 2010). Genomes that are subject to intense genetic drift are expected to be less well-optimized compared to those experiencing lower genetic drift. Michael Lynch proposed that variation in the ability to purge slightly deleterious mutations (*i*.*e*. variation in *N* _e_) can account for differences in genome characteristics among species (Lynch and Conery, 2003). This hypothesis has been empirically validated for multiple genome characteristics and phylogenetic clades. For example, it was shown that the genomes of crustacean species with low *N* _e_ values are larger than those of their sister species (Lefébure *et al*., 2017). Moreover, species with large *N* _e_ tend to have a lower mutation rate than species with low *N* _e_, illustrating the notion that natural selection acts to improve replication fidelity, within the constraints defined by random genetic drift (Lynch *et al*., 2016).

We recently examined the variations in transcriptome complexity across animal species in light of the “drift barrier” hypothesis (Bénitìere *et al*., 2024). In multicellular eukaryotes, the vast majority of genes give rise to multiple isoforms through alternative splicing (Chen *et al*., 2014). This phenomenon has attracted a great deal of interest since its discovery almost 50 years ago (Berget *et al*., 1977). Alternative splicing is commonly hypothesized to be adaptive, because it can increase the number of biological functions that are encoded in each genome. Indeed, numerous instances of alternative splicing patterns with beneficial effects have been identified (Mudge *et al*., 2011; Barbosa-Morais *et al*., 2012; Merkin *et al*., 2012; Reyes *et al*., 2013; Verta and Jacobs, 2022; Singh and Ahi, 2022; Wright *et al*., 2022). However, these examples represent only a small fraction of all splice variants that are now known, especially given the substantial detection power brought by next-generation RNA sequencing (RNA-seq) techniques. Many of the splice variants that can now be detected with RNA-seq are present at very low frequencies (Gonzàlez-Porta *et al*., 2013; Tress *et al*., 2017) and are poorly conserved during evolution (Barbosa-Morais *et al*., 2012; Merkin *et al*., 2012). It was thus hypothesized that they may be the result of errors of the splicing machinery, rather than functional isoforms (Pickrell *et al*., 2010; Gout *et al*., 2013; Xu and Zhang, 2014; Saudemont *et al*., 2017; Xu and Zhang, 2018; Liu and Zhang, 2018b,a; Xu *et al*., 2019; Xu and Zhang, 2020; Zhang and Xu, 2022). Notably, according to the “drift barrier” hypothesis, one may hypothesize that if alternative splicing (AS) primarily serves functional roles, the rate of alternative splicing should increase with *N* _e_. Conversely, if AS predominantly involves deleterious processes, its rate should decline with increasing *N* _e_. We applied this reasoning in our previous work (Bénitìere *et al*., 2024), which led us to deduce that AS is predominantly non-functional.

This methodology for exploring the impact of *N* _e_ on biological processes holds potential for broader applications. For example, one could examine the functional importance of alternative polyadenylation sites (Xu and Zhang, 2018). Such investigations demand cross-species comparative transcriptomics analyses, a task facilitated by the abundant availability of publicly accessible RNA-seq data. Yet, analysis of transcriptome sequencing data is resource-intensive in terms of time, energy, and computational power. To facilitate future analyses, we provide a comprehensive database that streamlines the process by offering pre-processed data. This dataset includes proxies for effective population size, sets of orthologous single-copy genes, gene expression levels, and intron-centered alternative splicing frequencies, along with phylogenetic trees to control for phylogenetic inertia. These resources have been compiled for 16,000 RNA-seq samples spanning 1,506 multicellular eukaryotic species.

This database, that we name GTDrift, complements other public transcriptomic data resources, such as Bgee (Bastian *et al*., 2020), which provides gene expression levels for 52 species (Version 15.0.1), but not alternative splicing frequencies. Other databases do provide alternative splicing frequencies. For example, MeDAS (Li *et al*., 2020) provides AS data for 18 metazoan species, and MetazExp (Liu *et al*., 2021) provides data for 72 metazoan species. This latter resource is substantial, including data for ∼ 53,000 RNA-seq samples. However, this database favors insects (53 species, with ∼ 26 000 RNA-seq samples for *Drosophila melanogaster*) and does not include any representative of the vertebrate clade, for which more computational resources are required because of their large genomes. Our database encompasses a broader phylogenetic distribution of species (Fig. 1), with 93 green plant species, 560 invertebrates and 853 vertebrates. Moreover, while other public databases such as MetazExp are aimed at biologists who want to analyze alternative splicing patterns in a gene-by-gene manner through web queries, in GTDrift we provide all data in flat files, which enable downstream computational analyses. GTDrift is thus mainly aimed at users with some computational skills. Nevertheless, we have created a user-friendly Shiny app to facilitate exploration of the database and species-specific data downloads (Chang *et al*., 2023).

**Figure 1.**
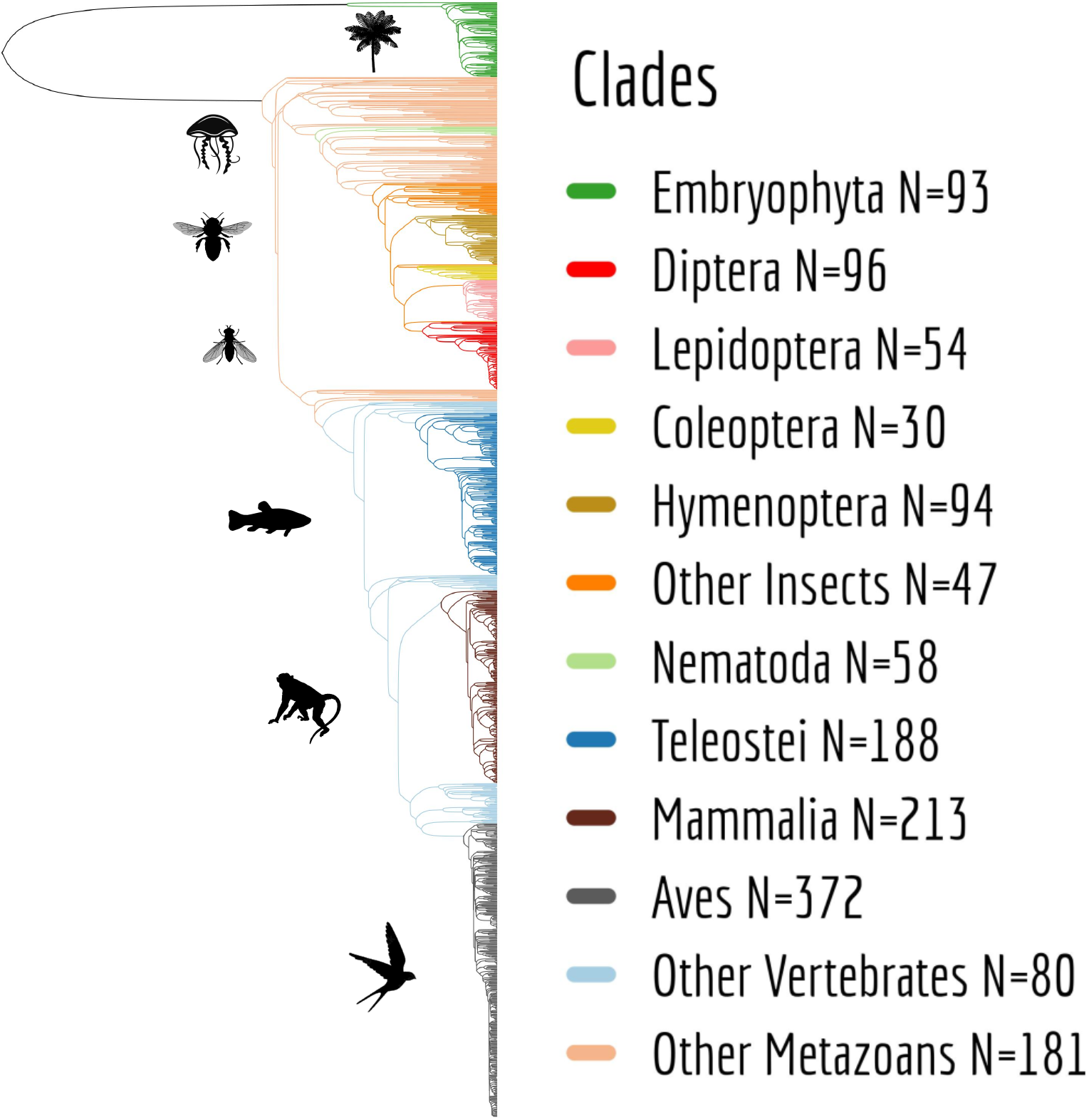
Phylogenetic distribution of the species included in the GTDrift database. The phylogeny was retrieved from TimeTree (Kumar *et al*., 2022). Not all species studied are present (N=1,220).

In GTDrift, we used assemblies and annotations data collected from The National Center for Biotechnology Information (NCBI) (Sayers *et al*., 2022), as well as publicly available RNA-seq data to investigate alternative splicing patterns and gene expression profiles. We computed summary statistics across all analyzed RNA-seq samples for each species, which enabled us to determine whether the available sequencing depth is sufficient for the study of alternative splicing. To ensure comparability across species, we annotated Benchmarking Universal Single Copy Orthologs (BUSCO) (Waterhouse *et al*., 2018) genes in all species and provide phylogenetic trees to control for phylogenetic inertia.

We believe that this tremendous amount of information should be shared with the scientific community, because it provides the means to investigate the impact of genetic drift on genome and transcriptome architecture, on a broad phylogenetic scale.

## Methods

### Species selection

The first criterion for species inclusion in GTDrift is the availability of a genome assembly and annotation in the NCBI database (NCBI Resource Coordinators, 2018; Sayers *et al*., 2022), as well as the availability of RNA-seq data in the Short Read Archive (Leinonen *et al*., 2011). We included 1,506 multicellular eukaryotic species. This collection encompasses 1,413 animal species as well as 93 species of green plants (Fig. 1). Our Snakemake pipeline can be applied to any species for which genome sequence, genome annotation and RNA-seq data are available, which will enable us to further expand GTDrift in the future(Mölder *et al*., 2021).

### Collecting life history traits

We queried several databases to acquire three specific life history traits, namely: maximum longevity, body mass, and body length. These traits were previously identified as suitable proxies for estimating the effective population size (Romiguier *et al*., 2014; Waples, 2016; Figuet *et al*., 2016; Galtier, 2016; Weyna and Romiguier, 2020). For eusocial species, which live in colonies and have both reproductive and non-breeding individuals, we gather data on the queen of the colony. For solitary species, we did not take into account the sex of the individuals, *i*.*e*. we retained the maximum value observed.

We employed several distinct methodologies to screen the databases. We initially used a manual approach to search across various sources of information, including scientific papers and databases.

We manually searched for information on life history traits from four prominent databases, which encompass diverse taxonomic groups. The Animal Ageing and Longevity Database (AnAge) (Tacutu *et al*., 2013), is renowned for its comprehensive collection of vertebrates, particularly mammals. The Encyclopedia of Life (EOL) (Wilson, 2003; Parr *et al*., 2014) encompasses a wide spectrum of species, prominently featuring invertebrates. The Animal Diversity Web (ADW) (Myers *et al*., 2023), is a particularly rich resource for invertebrates. The FishBase (Froese and Pauly, 2023) predominantly houses data on teleostei species. While AnAge furnishes extensive information regarding body mass and lifespan, it is lacking data pertaining to body length (Fig. 2A,B,C). Furthermore, as previously noted, certain databases are tailored to specific clades. For instance, in comparison to EOL and ADW, AnAge contains relatively fewer records for invertebrates (Fig. 2D,E,F).

**Figure 2.**
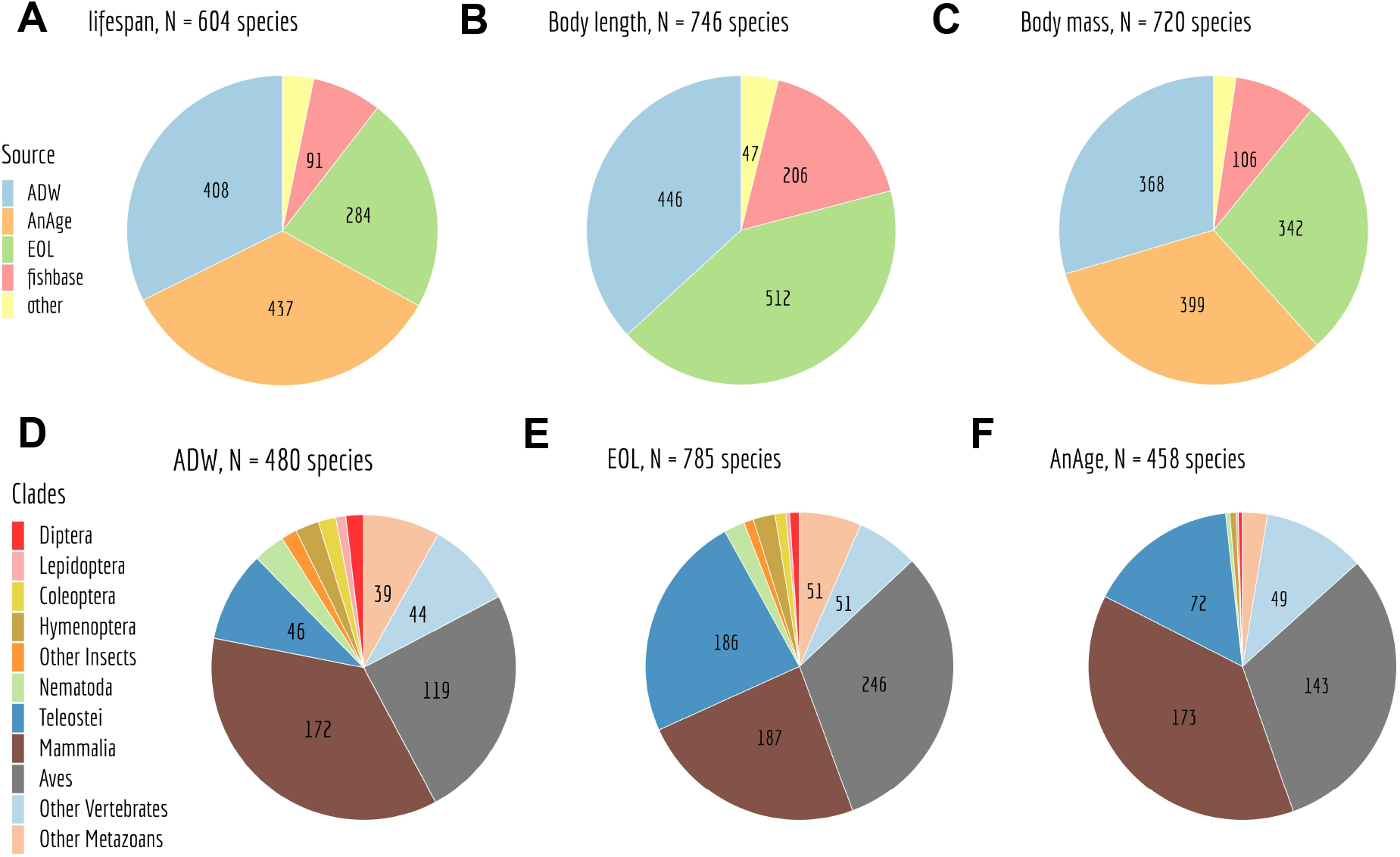
Representation of life history traits retrieved from diverse data sources. Depiction of data origins for lifespan (**A**), body length (**B**), and body mass (**C**). Additionally, distribution of species and their respective clades with at least one recorded life history trait in ADW (**D**), EOL (**E**), and AnAge (**F**).

We then made efforts to automate the manual search procedures. The primary automated procedure involved the development of a bash script, which utilized the Latin nomenclature of the species to navigate the textual content within the research pages of the 4 databases listed above. The bash script was designed to extract sentences, words, and numerical data in proximity to keywords such as “longevity”, “mass”, “weight”, and “length”, serving as indicators of relevant information. Its output was then reformatted through an R script. While this approach proved effective for databases like AnAge, EOL, and FishBase, its applicability to the ADW database was limited due to the manner in which information is embedded within textual paragraphs. Consequently, we employed an alternative method for the ADW database, involving Machine Learning and Natural Language Processing Question-Answering techniques. We obtained a trained model named “tinyroberta-squad2” from huggingface.co (Wolf *et al*., 2020). This model was used to answer questions related to specific attributes, such as ‘what is the body length ?’; ‘what is the body mass ?’; ‘what is the longevity ?’. Each question retrieved a pool of 100 potential answers derived from the database’s textual content, ranked by their predictive scores provided by the model.

We implemented an iterative selection process to identify the highest predicted answer containing relevant units and numeric values. To avoid redundancy, the selected answer was then removed from the text, and the process was repeated up to 10 times. The entire procedure was implemented in a Python script. We processed the script’s output to restructure the obtained results.

Discrepancies between the manual approach and the other two methodologies were further re-investigated manually and corrected as needed after a further re-reading of the text. As a result, the curated dataset that we share reflects our highest level of confidence.

In total, our data collection effort resulted in the acquisition of life history traits for 979 metazoan species.

### Acquisition of the reference genome sequence and annotations

Using the sra-tools software, we performed an automated identification of the reference genome for each species. Subsequently, we downloaded the annotation data in GFF format, the nucleotide coding sequences in FASTA format, and the peptide sequences in FASTA format from the NCBI database (Sayers *et al*., 2022).

### *dN/dS* pipeline

We developed a pipeline to estimate the rate of non-synonymous substitutions divided by the synonymous substitutions rate (*dN/dS*), representing the frequency at which non-synonymous changes occur relative to synonymous ones. Since non-synonymous substitutions are commonly perceived as errors, *dN/dS* serves as a measure of the rate of erroneous substitutions per neutral substitution. This ratio is directly dependant of *N* _e_ as it is jointly determined by the distribution of selection coefficient of new mutations (*s*) and the magnitude of genetic drift as defined by *N* _e_ (Yang and Nielsen, 1998; Nielsen and Yang, 2003). The transcriptome-wide *dN/dS* is expected to rise over prolonged periods of small *N* _e_ due to the increasing number of slightly deleterious mutations reaching fixation (Ohta, 1992; Galtier, 2016).

Estimating the *dN/dS* necessitates the annotation of genes shared across all species, their evolutionary history depicted by a phylogenetic tree, and finally a comparative analysis of site evolution to derive the *dN/dS* ratio.

### BUSCO genes identification

We used the BUSCO v.3.1.0 software to identify single-copy orthologous genes within three datasets selected from OrthoDB v9 (Zdobnov *et al*., 2017): eukaryota (N=303 genes), embryophyta (N=1,440 genes) and metazoan (N=978 genes) sourced from BUSCOv3 (Waterhouse *et al*., 2018; Seppey *et al*., 2019; Manni *et al*., 2021). The search was performed against the longest annotated protein sequences *per* gene within each genome.

### Phylogenetic tree reconstruction

Due to the considerable time and resource demands associated with phylogenetic inference for large numbers of species, we employed a strategy in which the analysis was partitioned by clades. On initial releases of the database, which did not encompass all current species, we performed 3 comparable and independent analyses that rely on the three BUSCO datasets, corresponding to the following lineages: eukaryota, embryophyta and metazoa. For each BUSCO dataset, we selected a subset of species that matched the lineage of interest from the available database records at the time of analysis. All of these selected species underwent transcriptomic analyses (see Transcriptomic analyses). We then collected the longest corresponding proteins identified in each species for each BUSCO gene family. We removed proteins for which the amino acid sequence provided with the annotations did not perfectly correspond to the translation of the corresponding coding sequences. We then aligned the resulting sets of protein-coding sequences for each BUSCO gene, using the codon alignment option in PRANK v.170427 (Löytynoja and Goldman, 2008). We translated the codon alignments into protein alignments using the R package seqinr (Charif and Lobry, 2007).

A filter was applied to retain only genes for which enough species have been detected (85% of the analyzed species), reducing the eukaryota set to 126 genes (embryophyta N=387 genes, metazoa N=731 genes). Then, species were removed from the analysis if they had less than 80% of the studied genes, reducing the number of studied species from 336 to 279 for the eukaryota BUSCO dataset (embryophyta 93 to 80 species, metazoa 293 to 257 species).

To infer the phylogenetic tree rapidly, we sub-sampled the resulting multiple alignments, selecting alignments with the highest number of species (eukaryota N=25 genes, embryophyta N=77 genes, metazoa N=146 genes). We then concatenated these alignments and kept sites that were aligned in most of the analyzed species (see information provided in the supplementary archive for more details). The final alignment for the eukaryota BUSCO dataset included 279 taxa (embryophyta N=80 species, metazoa N=257 species) taxa and 600,807 sites (embryophyta N=670,083 sites, metazoa N=3,135,111 sites). We used RAxML-NG (Kozlov *et al*., 2019), to infer the species phylogeny on these final alignments. RAxML was set to perform one model *per* gene with a fixed empirical substitution matrix (LG), empirical amino acid frequencies from alignment (F) and 8 discrete GAMMA categories (G8). These parameters were specified in a partition file with one line *per* BUSCO gene multiple alignment. The analysis generated at least 10 starting trees. The best-scoring topology was kept as the final ML tree and 10 bootstrap replicates have been generated.

The phylogenetic trees were rooted using as a reference source the TimeTree phylogeny, which synthesizes data from numerous published studies, despite its incomplete representation of all species (Kumar *et al*., 2022).

To encompass a broader spectrum of the species included in our latest database release, the one published here, we also reconstructed phylogenetic trees *per* clade. To do this, we divided the full set of metazoan species in 9 groups (Hymenoptera, Diptera grouped with Lepidoptera under the superorder Mecopterida, Nematoda, other insects, Aves, Mammalia, Teleostei, other vertebrates, and finally other invertebrates). We ranked the 731 metazoan BUSCO genes in decreasing order of the number of species in which they were annotated. We then selected as a basis for the analyses the 73 genes at the top of this list, corresponding to the top 10% genes. We applied the protocol described above to each individual clade. The resulting clade-specific trees were merged using outgroup species as a reference point to construct the complete metazoan phylogenetic tree.

### *dN /dS* computation

We computed *dN/dS* ratios for BUSCO gene families that were present in at least 85 percent of the species under investigation. We conducted four independent analyses. We first analyzed each of the three BUSCO gene sets: eukaryota (N=126 genes), embryophyta (N=387 genes), metazoa (N=731 genes). We also performed an analysis ‘*per* clade’, as explained above for the phylogenetic tree reconstruction, using the same 731 genes preselected in the metazoa analysis. Codon alignments obtained using PRANK (Löytynoja and Goldman, 2008) served as the basis for this estimation. To manage the computational memory demands during the substitution rate estimation step, we segmented the sequence alignments into clusters. Following the approach recommended by Bolívar *et al*. (2019), these clusters were defined based on the average GC3 content across species, in order to group genes with similar parameters. We then concatenated the alignments within each group, obtaining alignments that were 200 kb long on average. This process yielded 13 groups for eukaryota (15 for embryophyta and 73 for metazoa). We used bio++ v.3.0.0 libraries (Dutheil and Boussau, 2008; Guéguen *et al*., 2013; Bolívar *et al*., 2019) to estimate the *dN/dS* on each branch of the phylogenetic tree, for each concatenated alignment.

In a first step, we used an homogeneous codon model implemented in bppml to infer the most likely branch lengths, codon frequencies at the root, and substitution model parameters. We used YN98 (F3X4) (Yang and Nielsen, 1998) substitution model, which allows for different nucleotide content dynamics across codon positions. In a second step, we used the MapNH substitution mapping method to count synonymous and non-synonymous substitutions (Dutheil *et al*., 2012; Guéguen and Duret, 2018). We defined *dN* as the total number of non-synonymous substitutions divided by the total number of non-synonymous mutational opportunities, both summed across concatenated alignments, for each branch of the phylogenetic tree.

Likewise, we defined *dS* as the total number of synonymous substitutions divided by the total number of synonymous mutational opportunities, both summed across concatenated alignments. The *per* -species *dN/dS* corresponds to the ratio between *dN* and *dS*, on the terminal branches of the phylogenetic tree. We also provide the *dN* and *dS* values for each branch within the phylogenetic trees.

For the ‘*per* clade’ approach, the results pertaining to distinct clades were combined in a single table.

### Transcriptomic analyses

We developed a pipeline facilitating the detection of alternative splicing events within genes. This process entails the selection of RNA-seq data, subsequent alignment to the reference genome, and the identification of splicing events through the recognition of introns. Utilizing the aligned transcriptomic data, we computed gene expression levels across each sample.

### Polymorphism-derived *N* _e_ estimates

In Lynch *et al*. (2023) the *per* species germline mutation rate (*µ*) and level of neutral diversity (*π*_*s*_) was integrated into the equation *N*_*e*_= *π*_*s*_*/*4*µ*, to produce what we named a “polymorphism-derived *N* _e_”. This more direct estimates of *N* _e_ was calculated for 65 of the species in our dataset.

Additionally, we expanded our dataset with the *N* _e_ estimate for *C. nigoni* by including *π*_*s*_ = 0.06 (Asher Cutter, personal communication) and *µ* = 1.3 *×* 10^−9^ (Denver *et al*. (2012); assuming a similar mutation rate as in *C. briggsae*).

### Selection of the RNA-seq samples

To extract RNA-seq data, we queried the Short Read Archive (SRA) database for samples where the library source was ‘TRANSCRIPTOMIC’ and the library strategy was ‘RNA-seq’.

For perfect comparability of transcriptome data among species, we would need to have the same representation of individual tissues, developmental stages *etc*. for each species, with data generated with the same protocol by the same person. However, such data exist only for limited sets of species (*e*.*g*., Cardoso-Moreira *et al*. (2019)). Here, we decided not to filter the RNA-seq samples on criteria pertaining to sample origin or experimental protocols, mainly because the relevant information is not always provided in sufficient detail in the SRA database (Leinonen *et al*., 2011). Moreover, depending on the clade, the biological sample of origin can vary from “whole body” in insects, to specific tissues or cell types in mammals. Thus, perfectly comparable sample collections are difficult to obtain across such a broad phylogenetic scale.

Rather than filtering samples on these criteria prior to inclusion in the database, in GTDrift we provide users with the information collected from SRA for all RNA-seq samples. This information includes the library type, the date of extraction and the name of the laboratory that performed the experiment (see Description of the data available in GTDrift).

After evaluating the amount of RNA-seq data that is needed to evaluate global alternative splicing patterns for each species (see below), we decided to include a maximum of 50 RNA-seq samples *per* species in GTDrift.

We included more than 50 samples for 150 species (43 embryophyta, 107 metazoa), for which we performed more detailed analyses, considering various tissues or developmental stages.

In the current version of GTDrift, the RNA-seq dataset encompasses a total of 491 distinct species, including 92 plants and 399 animals. (Fig. 3A).

**Figure 3.**
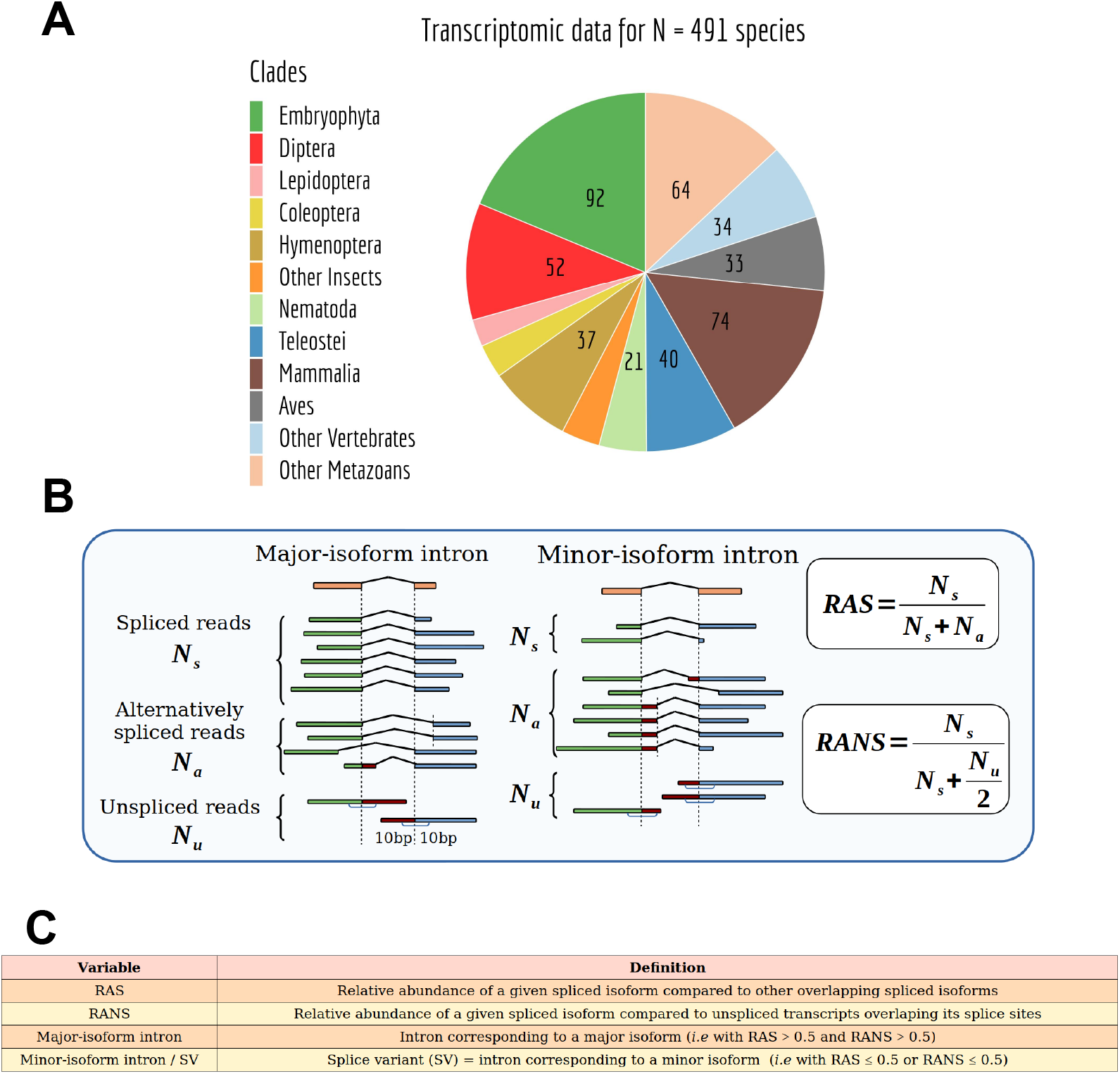
Species with transcriptomic data and alternative splicing estimation. (*cf* Fig. 2A Bénitìere *et al*. (2024)) **A**: Taxonomic distribution of the species for which transcriptomic data was included in GTDrift. **B**: Definition of the variables used to compute the relative splicing frequency of a focal intron, compared to splice variants with a common alternative splice boundary (RAS) or compared to the unspliced form (RANS): N_s_: number of spliced reads corresponding to the precise excision of the focal intron; N_a_: number of reads corresponding to alternative splice variants relative to this intron (*i*.*e*. sharing only one of the two intron boundaries); N_u_: number of unspliced reads, co-linear with the genomic sequence. **C**: Definitions of the main variables used in this study. The definition of the variables corresponds to the one provided in Bénitìere *et al*. (2024).

### Indexing genomes and aligning RNA-seq data

The RNA-seq alignment phase represents the most time-consuming stage in the pipeline (Fig. 4), and can extend up to one week when utilizing 16 cores for each RNA-seq dataset, particularly for larger genomes such as those of mammals.

**Figure 4.**
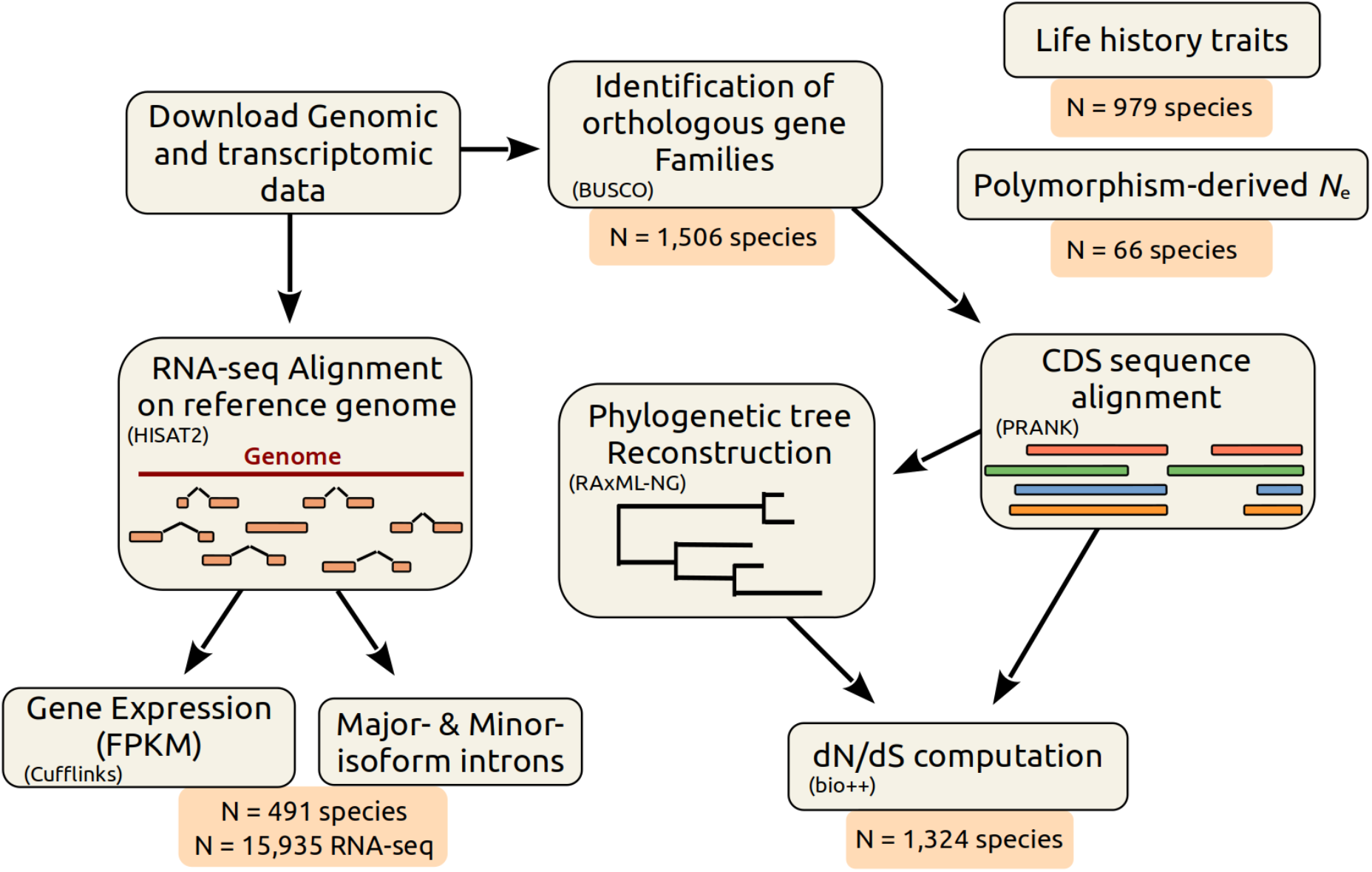
Description of the bioinformatic analysis pipeline. (Adapted from Supplementary Fig. 11 Bénitìere *et al*. (2024)) First, we retrieved genomic sequences and annotations from the NCBI Genomes database. We aligned RNA-seq reads on the corresponding reference genomes with HISAT2. We used these alignments to estimate various variables related to splicing patterns (see Fig. 2), to compute the AS rate, and to estimate gene expression using Cufflinks. To compute the *dN /dS* ratios, we first identified BUSCO genes with BUSCOv3 and aligned their coding sequences (CDS) using PRANK (codon model). We reconstructed a phylogenetic tree using RAxML-NG. Using bio++, we estimated *dN /dS* along the phylogenetic tree on concatenated alignments. This pipeline was previously used in Bénitìere *et al*. (2024).

For this step, HISAT2 version 2.1.0 was employed to align RNA-seq reads to the respective reference genomes (Kim *et al*., 2019). To enhance the sensitivity of splice junction detection, we constructed genome indexes incorporating annotated intron and exon coordinates along with genome sequences. The maximum permitted intron length was set at 2,000,000 base pairs. The processed and compressed files generated during this procedure can amass a size exceeding 20 terabytes.

We extracted intron coordinates from the HISAT2 alignments, utilizing a custom Perl script that scanned for CIGAR strings containing “N” characters, which indicate skipped regions in the reference sequence. For intron identification and quantification, we exclusively utilized uniquely mapped reads with a maximum mismatch fraction of 0.02. In the context of new intron identification, we imposed a minimum anchor length (*i*.*e*., part of the read that spans each of the two exons flanking a given intron) of 8 base pairs. We then quantified intron splicing frequencies by including aligned reads with a minimum anchor length of 5 base pairs. We retained predicted introns exhibiting GT-AG, GC-AG, or AT-AC splice signals and determined the intron strand based on the splice signal.

Introns were assigned to genes if at least one of their boundaries was within 1 base pair of annotated exon coordinates, combined across all isoforms for each gene. Intron assignments were limited to those that could be unambiguously associated with a single gene. Notably, we differentiated between annotated introns, present in the reference genome annotations, and unannotated introns, identified through RNA-seq data and assigned to previously annotated genes.

We identified introns situated within protein-coding regions. To do this, for each protein-coding gene, we extracted annotated start and stop codon positions across all annotated isoforms. The minimum start codon and maximum end codon positions were identified, and introns located upstream or downstream of these extreme coordinates were considered as interrupting untranslated regions.

### Alternative splicing variables

For each intron, we recorded two key variables: N_s_ representing the number of reads corresponding to the precise removal of the intron (referred to as spliced reads), and N_a_ representing the count of reads supporting alternative splicing events (*i*.*e*. spliced variants sharing only one of the two boundaries of the focal intron). Additionally, we denoted N_u_ as the count of unspliced reads that align linearly with the genomic sequence and span at least 10 base pairs on both sides of an exon-intron junction. These definitions are visually clarified in (Fig. 3B,C). Subsequently, we introduced the relative measurement of the target intron’s abundance compared to introns with a single alternative splice boundary 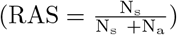, as well as relative to unspliced reads 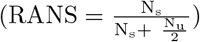.

To compute these ratios, we required at least 10 reads in their denominators. Thus, we computed the RAS only when (N_s_ + N_a_) ≥ 10, and the RANS only when 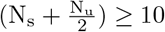. We divided N_u_ by 2 because unspliced reads that span the two intron boundaries likely refer to the same intron retention event. If these conditions were not met, the resulting values were designated as unavailable (NA). These ratios were computed utilizing data from all available RNA-seq samples, unless explicitly specified (*e*.*g*. in sub-sampling analyses). Based on these ratios, we divided introns into three categories: major-isoform introns, defined as those introns that have RANS *>* 0.5 and RAS *>* 0.5 (these likely correspond to the introns of major isoforms (Gonzàlez-Porta *et al*., 2013; Tress *et al*., 2017; Bénitìere *et al*., 2024); minor-isoform introns, defined as those introns that have RANS ≤ 0.5 or RAS ≤ 0.5 (these introns are detected in a minority of transcripts); unclassified introns, which do not satisfy the above conditions.

### Gene expression estimation

Gene expression levels were computed using Cufflinks version 2.2.1 (Trapnell *et al*., 2010; Roberts *et al*., 2011), utilizing the read alignments obtained with HISAT2 for each individual RNA-seq sample. We thus evaluated gene expression levels with the Fragment *Per* Kilobase of exon *per* Million mapped reads (FPKM) method. To determine the representative expression level of each gene, the mean FPKM was calculated across all samples, taking into consideration the sequencing depth of each sample, called ‘weighted FPKM’. We used this measure to evaluate the relationship between alternative splicing rates and gene expression levels, within each species.

### Estimation of the sequencing depth

We determined for each gene the union of all annotated exon coordinates (termed here exon blocks). Using bedtools v2.25.0 (Quinlan and Hall, 2010), we assessed the read coverage at each position of the exon blocks. The average exonic *per* -base read coverage was subsequently computed for each gene. The sequencing depth of a given sample was evaluated through the median *per*-base read coverage across BUSCO (Benchmarking Universal Single-Copy Orthologs) genes.

### Data visualisation using a Shiny app

A Shiny app available at https://lbbe-shiny.univ-lyon1.fr/ShinyApp-GTDrift/ was deployed to allow users to visualize and compare the summarized data (Chang *et al*., 2023). Most of the graphics shown in this paper are directly reproducible from the app. In this app, users can also visualize and download intra-species variables, for example comparing introns or gene characteristics. Furthermore, a specific tab is dedicated to the investigation of gene structure in relation to the splicing attributes found in the underlying database. Users can also visualize the phylogenetic tree and employ these trees for conducting Phylogenetic Generalized Least Square regression analyses.

The app is organized in several panels or “tabs” in the web page.

The tab ‘Inter-species graphics’ facilitates the comparison of genome characteristics across different species through graphical representation. Additionally, users have the option to upload their own data in a tab-separated text format, where each species is represented in a separate row, with the variables of interest organized in columns. An example of such a tabular dataset can be found in the repository of the Shiny app.

The ‘Inter-species Axis’ tab explains the variables available in the ‘Inter-species graphics’ tab.

The ‘Intra-species graphics’ tab permits the exploration of characteristics within a species, focusing on introns or on genes. Furthermore, users have the ability to download metadata related to BUSCO annotation, gene expression profile, or intron splicing events (see Methods).

The ‘Intra-species Axis’ tab describes the variables featured in the ‘Intra-species graphics’ tab.

Within the ‘Gene structure’ tab, users can delve into the introns detected in RNA-seq alignments for a specific gene. These introns are color-coded based on various criteria, including their location within the CDS or outside of it, as well as whether they are classified as major or minor-isoform introns (see Methods).

The ‘Phylogenetic tree’ tab facilitates the examination of phylogenetic trees used for conducting Phylogenetic Generalized Least Squares regression within the ‘Inter-species graphics’ tab.

## Data and code availability

The database is provided on Zenodo with the DOI: https://doi.org/10.5281/zenodo.10017653.

All processed data that we generated and used in this study, as well as the scripts that we used to analyze the data and to generate the figures, are available at the following Zenodo DOI: https://doi.org/10.5281/zenodo.10022493. Finally, the Shiny app is available at: https://lbbe-shiny.univ-lyon1.fr/ShinyApp-GTDrift/ and on Zenodo with the DOI: https://doi.org/10.5281/zenodo.10022520.

## Results

### Description of the data available in GTDrift

In GTDrift, we provide a manageable number of compressed data tables for each species processed via our pipeline (Fig. 4). Tables are stored in tab-delimited text format, which makes them easy to access for users with experience in bioinformatics. They are user-friendly because of the simplicity of their contents. To access these tables, users can visit the Zenodo DOI: https://doi.org/10.5281/zenodo.10017653 and select their desired data type. The data can also be easily explored through a web application written in Shiny at https://lbbe-shiny.univ-lyon1.fr/ShinyApp-GTDrift/. Data exploration is thus easily accessible even for users who do not have a background in bioinformatics.

Our database is centered around transcriptomics data. At the time of publication, the database contained over 15,935 RNA-seq samples distributed over 491 embryophytes and metazoans (Fig. 1), providing gene expression and alternative splicing events data. Additionally, we have enriched the database with annotations for orthologous single-copy genes (BUSCO genes) and proxies of effective population size, including the molecular evolutionary rate *dN/dS*, the polymorphism-derived *N* _e_ estimates and life history traits such as longevity, body mass, body length. We used similar types of data in our recent publication exploring the relationship between random genetic drift and alternative splicing patterns (Bénitìere *et al*., 2024). However, here we provide considerably more data, for 1,506 species compared to 53 in this publication.

Below, we provide information on the data types that are currently available in GTDrift for the species listed in the table labeled ‘list species.tab’. This table contains additional information, such as genome/annotation assembly accession, number of RNA-seq samples for each studied species, species taxonomy, *etc*.

### Life history traits and polymorphism-derived *N* _e_

The table labeled ‘life history traits and polymorphism derived Ne.tab’ comprises values pertaining to three distinct traits (body mass, longevity, and body length), for 979 species. This table includes bibliographic references which attribute these values to each species. The species are defined by their scientific names and by the corresponding NCBI taxonomy identifier (taxID). Additionally, this table contains polymorphism-derived *N* _e_ estimates for 66 species.

### Protein-coding sequence evolution features

We provide estimates of the representative *dN/dS* ratio for most species (N=1,324 species after filtering for a sufficient number of annotated orthologous genes). The data are available in the directory ‘dNdS’.

We provide the phylogenetic tree of the studied species, with the *dN/dS* ratios as branch lengths, in the Newick file format. We provide this data separately for the four approaches used to estimate the ratios *dN/dS*, using the eukaryota, embryophyta or metazoa BUSCO gene sets, or a different gene set for each clade (see Methods). In addition, we provide a table comprising the *dN* and *dS* values for each terminal branch of the phylogenetic tree, along with the species scientific name and NCBI taxonomy ID, for each of the four approaches.

### Gene expression

In the ‘Transcriptomic’ directory, each species is represented by a dedicated table named ‘by gene analysis.tab.gz’. This table contains annotated gene coordinates, the mean and median FPKM (Fragments *Per* Kilobase of exon *per* Million mapped reads) across samples. Additionally, the table includes information about RNA-seq read coverage for exonic regions for each gene, including the total read coverage across samples. The individual gene expression data for each RNA-seq experiment can be accessed within the ‘RUN’ directory. The data are provided in a separate directory for each SRA accession number. The file ‘by gene db.tab.gz’ containing the exon coverage and the FPKM measured for each gene corresponding in line to the previous file ‘by_gene_analysis.tab.gz’.

### Alternative splicing data

For each species, we provide a summarized table named ‘by intron analysis.tab.gz’, containing for each intron the cumulative counts of spliced reads (N_s_), the number of reads supporting alternative splicing of this introns (N_a_), and the number of unspliced reads overlapping with this intron (N_u_) detected through RNA-seq analysis (see Methods). This table contains data combined across all analyzed RNA-seq samples. Detailed information for individual RNA-seq experiments can be found within the ‘RUN’ directory, in the file ‘by intron db.tab.gz’. In these files, introns are listed in the same order as in the file ‘by_intron_analysis.tab.gz’.

### RNA-seq sample description

In the file named ‘SRAruninfo.tab’, we provide information extracted from the SRA database, for each RNA-seq sample. Depending on the sample, this information can include the library source, the tissue from which the sample is derived, the sex of the sampled individual, the lab that conducted the analysis, the methods used to prepare the library, *etc*.

### BUSCO gene identification

In the directory ‘BUSCO annotations’, we provide the correspondence between NCBI gene identifiers and BUSCO gene identifiers, determined for three distinct BUSCO datasets: eukaryota, metazoa, and embryophyta.

### Data quality validation

#### Acquiring life history traits

To facilitate the acquisition of life history traits, we have devised and shared a pipeline that uses an automatic screening technique complemented by a Machine Learning method.

To assess the effectiveness of the automatic screening technique that we used to extract life history traits from various databases, we conducted a comparative analysis, contrasting it with the manual methodology. We also compared it to the Machine Learning (ML) approach for the ADW database. The screening procedure yielded accurate information with varying false positive rates depending on the source database, as follows: AnAge (98.9% accuracy; 0% false positive), fishbase (100%; 0.2%), EOL (94.5%; 0.2%), and ADW (87.9%; 5.4%). These results highlight the utility of our screening pipeline for identifying three key life history traits across AnAge, EOL, ADW, and fishbase databases.

For the ADW database, the ML approach exhibited a slight advantage over the screening method, and its results did not completely align with those obtained through the screening approach. Specifically, for life history traits, the ML approach correctly retrieved 89.8% of the results obtained through the manual approach, while introducing a 9.2% false positive rate.

When combining both the ML approach and the screening process, we achieved a 95.1% accuracy rate in identifying positive cases. However, a 7.3% error rate persisted in this merged approach.

In GTDrift, we provide data corresponding to a synthesis of the three methodologies including only manually-checked values (see Methods).

#### Estimating the intensity of random drift

As expected, a positive correlation is observed in Fig. 5A,B between the different life history traits, used as indirect predictors of the effective population size (*N* _e_) (Romiguier *et al*., 2014; Waples, 2016; Figuet *et al*., 2016; Galtier, 2016; Weyna and Romiguier, 2020).

**Figure 5.**
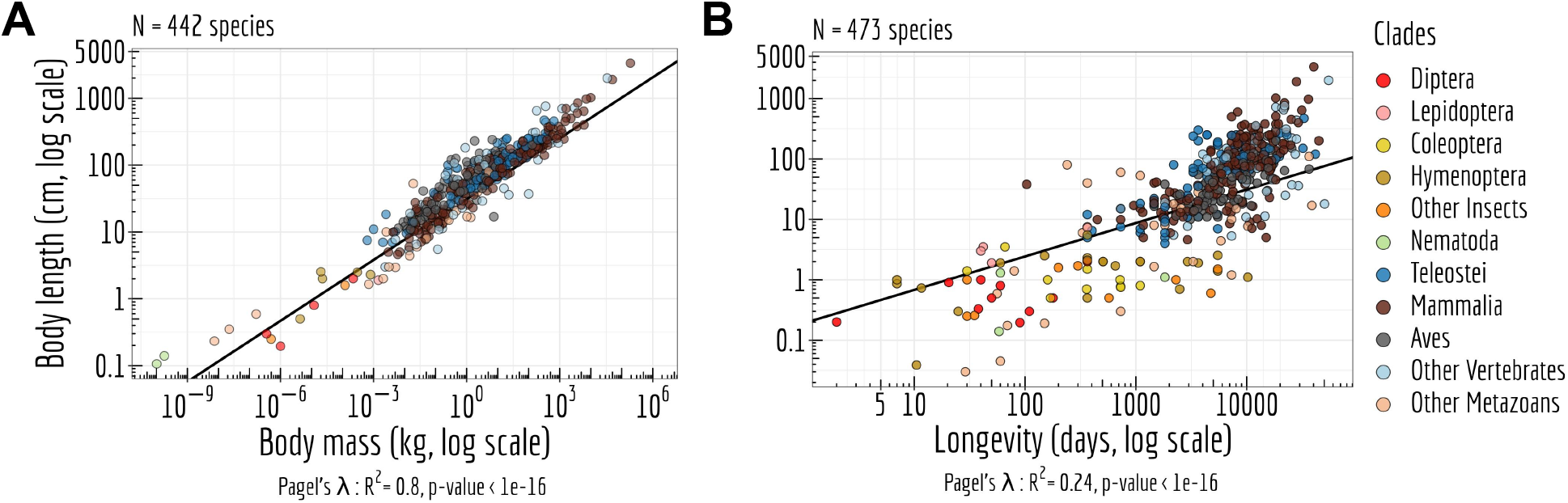
*N* _e_ proxies. **A**: Relationship between body length (cm, log scale) and the body mass (kg, log scale). **B**: Relationship between body length (cm, log scale) and longevity (days, log scale) of the organism. Each dot represents one species (colored by clade). **A**,**B**: Pagel’s *lambda* model is used to take into account the phylogenetic structure of the data in a regression model.

When examining the *dN/dS* ratio across distinct time scales and using various BUSCO datasets, we consistently observe comparable *dN/dS* ratios at terminal branches. This uniformity across a range of methodological approaches highlights their concordance (Fig. 6A,B).

**Figure 6.**
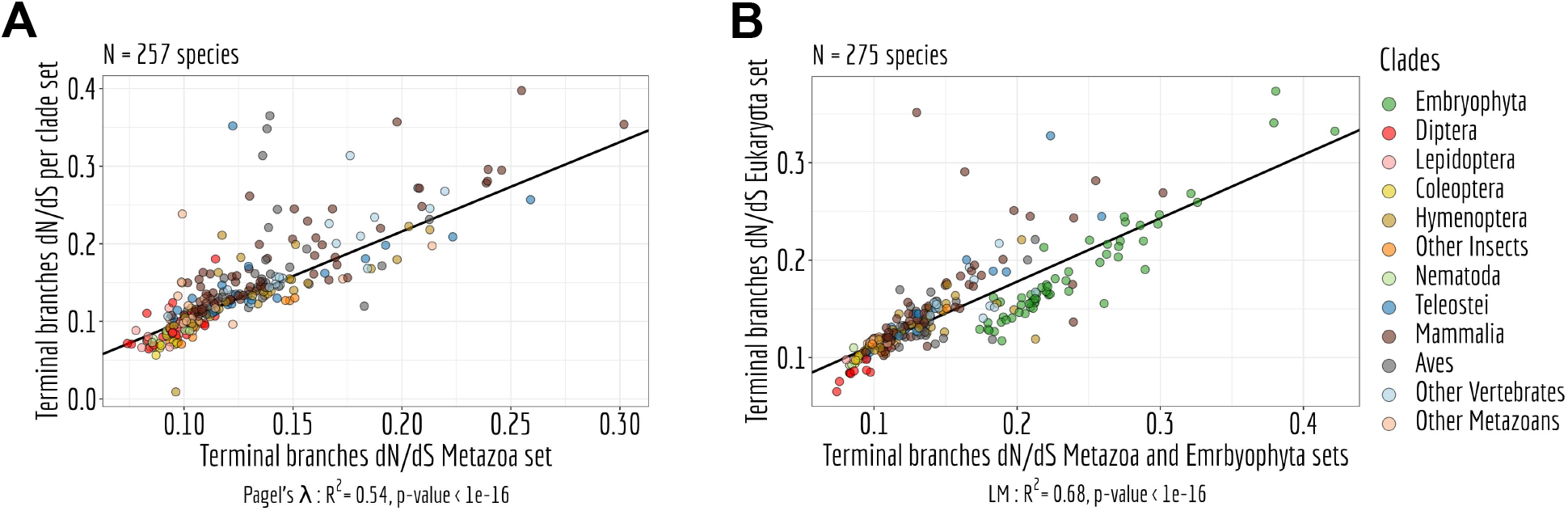
Reproducibility of the *dN /dS* ratio. **A**: Relation between the *dN /dS* ratio on terminal branches of the phylogenetic tree of the metazoa set compared to the ones measured in the *per* clades set. **B**: Relation between the *dN /dS* ratio on terminal branches of the phylogenetic tree of the eukaryota set compared to the ones measured in the embryophyta and the metazoa set. **A**,**B**: LM stands for Linear regression Model and Pagel’s *lambda* model is used to take into account the phylogenetic structure of the data in a regression model.

Furthermore, all the above proxies of *N* _e_ (*i*.*e*. longevity, body mass, body length and *dN/dS*) significantly correlate with a more direct *N* _e_ proxy, *i*.*e*. the polymorphism-derived *N* _e_ (Fig. 7; see Lynch *et al*. (2023)).

**Figure 7.**
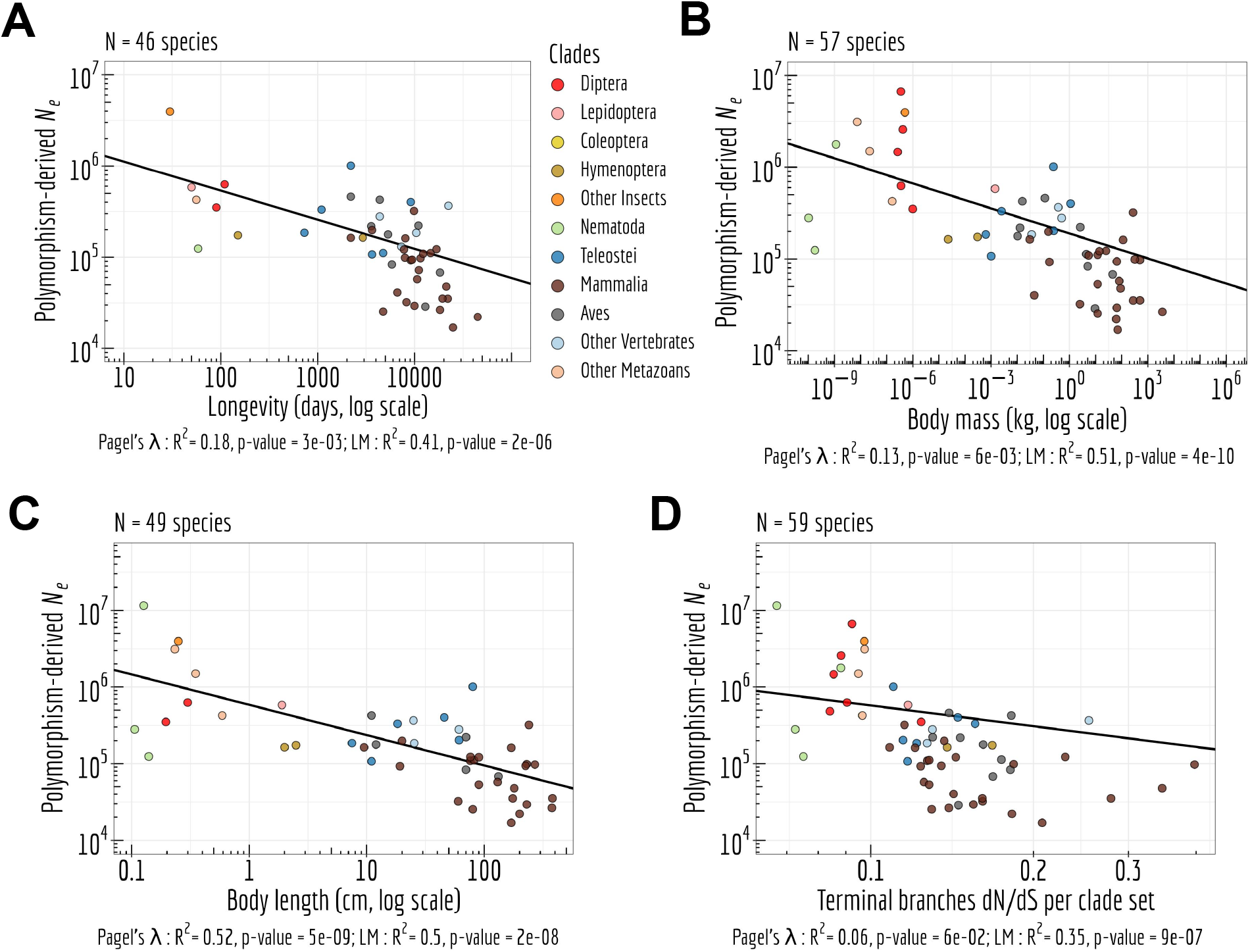
Interplay between *N* _e_ proxies. Correlation between the polymorphism-derived *N* _e_ and four other, more indirect, proxies of *N* _e_: life history traits such as longevity (days, log scale) (**A**), body mass (kg, log scale) (**B**), body length (cm, log scale) (**C**), and the *dN /dS* ratio on terminal branches of the phylogenetic tree of the *per* clade set (**D**). Pagel’s *lambda* model is used to take into account the phylogenetic structure of the data in a regression model.

### Quality of genome annotations

To assess gene expression levels and alternative splicing patterns, the quality of genome annotations is of paramount importance. We evaluated genome annotation quality by examining the presence of BUSCO genes. We note that the results depend on the BUSCO dataset that is used as a starting point. When using the BUSCO dataset designed for eukaryota, which comprises 303 genes, we have effectively identified nearly all single-copy orthologous genes, and this feature exhibits a high degree of homogeneity across different species (Fig. 8). However, the aves clade demonstrates a deficiency in the number of BUSCO genes compared to the anticipated count based on BUSCO expectations. This is expected given the known genome incompleteness problem for this clade, due to the presence of GC-rich chromosomes (Li *et al*., 2022).

**Figure 8.**
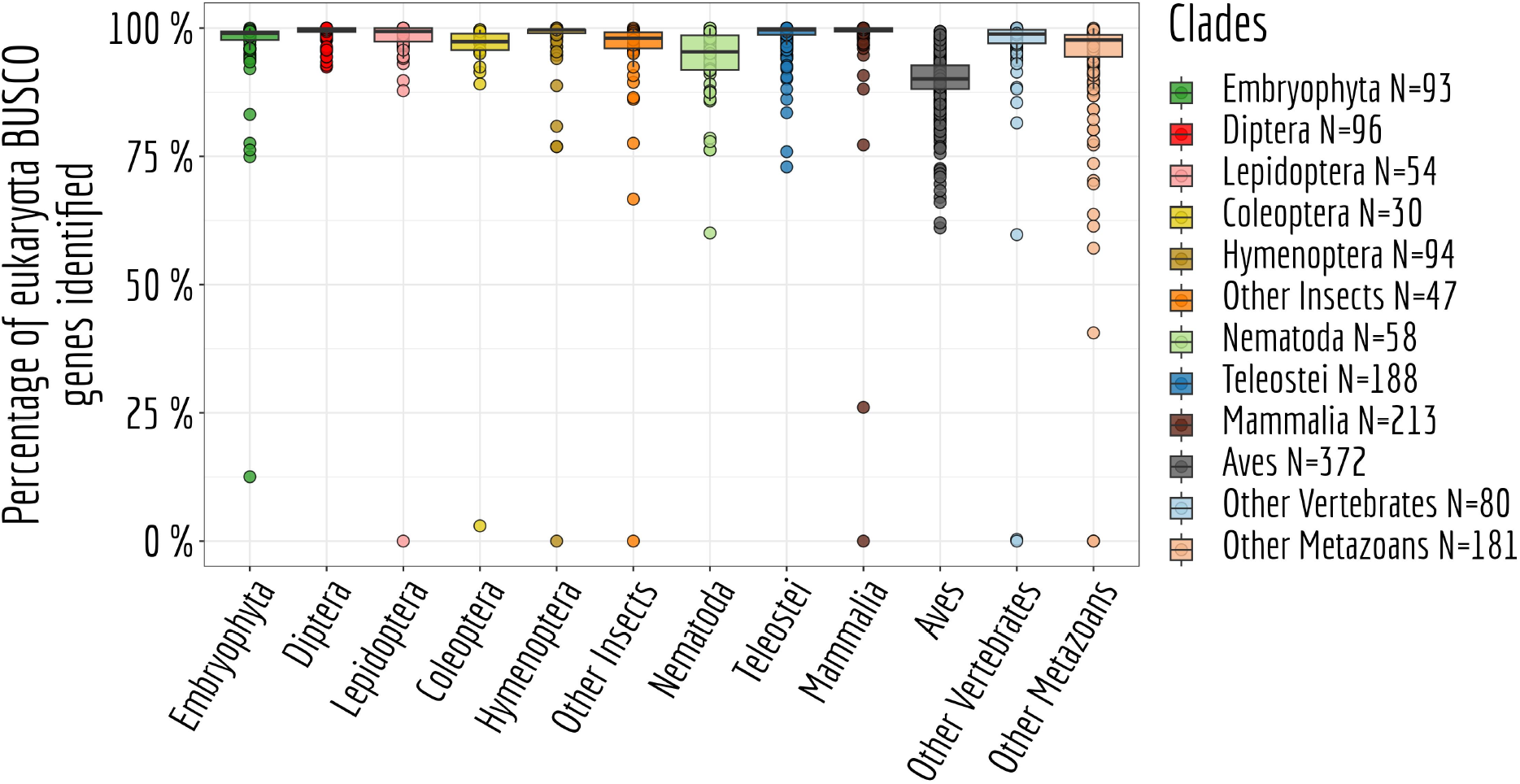
BUSCO genes annotation. Proportion of BUSCO genes, from the BUSCO gene set eukaryota (N=303 genes), identified in each species.

Because the eukaryota BUSCO gene set is limited, we also performed gene identification for the metazoa and embryophyta BUSCO datasets, leading to substantially larger collections of genes. Specifically, we detected 978 BUSCO genes for the metazoa dataset and 1,440 genes for the embryophyta dataset.

### Spliced introns classification

A significant body of literature has consistently reported that the majority of genes typically exhibit one predominant isoform (Gonzàlez-Porta *et al*., 2013; Tress *et al*., 2017). This isoform is commonly termed “major isoform”. Here, we aimed to assess the influence of sequencing depth on the identification of major-isoform introns, that is, those introns that belong to major isoforms (see Alternative splicing variables). Employing the model organism *Drosophila melanogaster*, we randomly selected between 1 and 20 RNA-seq samples.

For each subset of samples, we computed the median read coverage across the exons of BUSCO genes, providing a standardized measure of transcriptome sequencing depth that can be compared across different species. Additionally, we tallied the count of introns falling into various categories (major-isoform introns, minor-isoform introns or unclassified introns - see Methods) for each subset of samples. This entire process was repeated 10 times (Fig. 9A).

**Figure 9.**
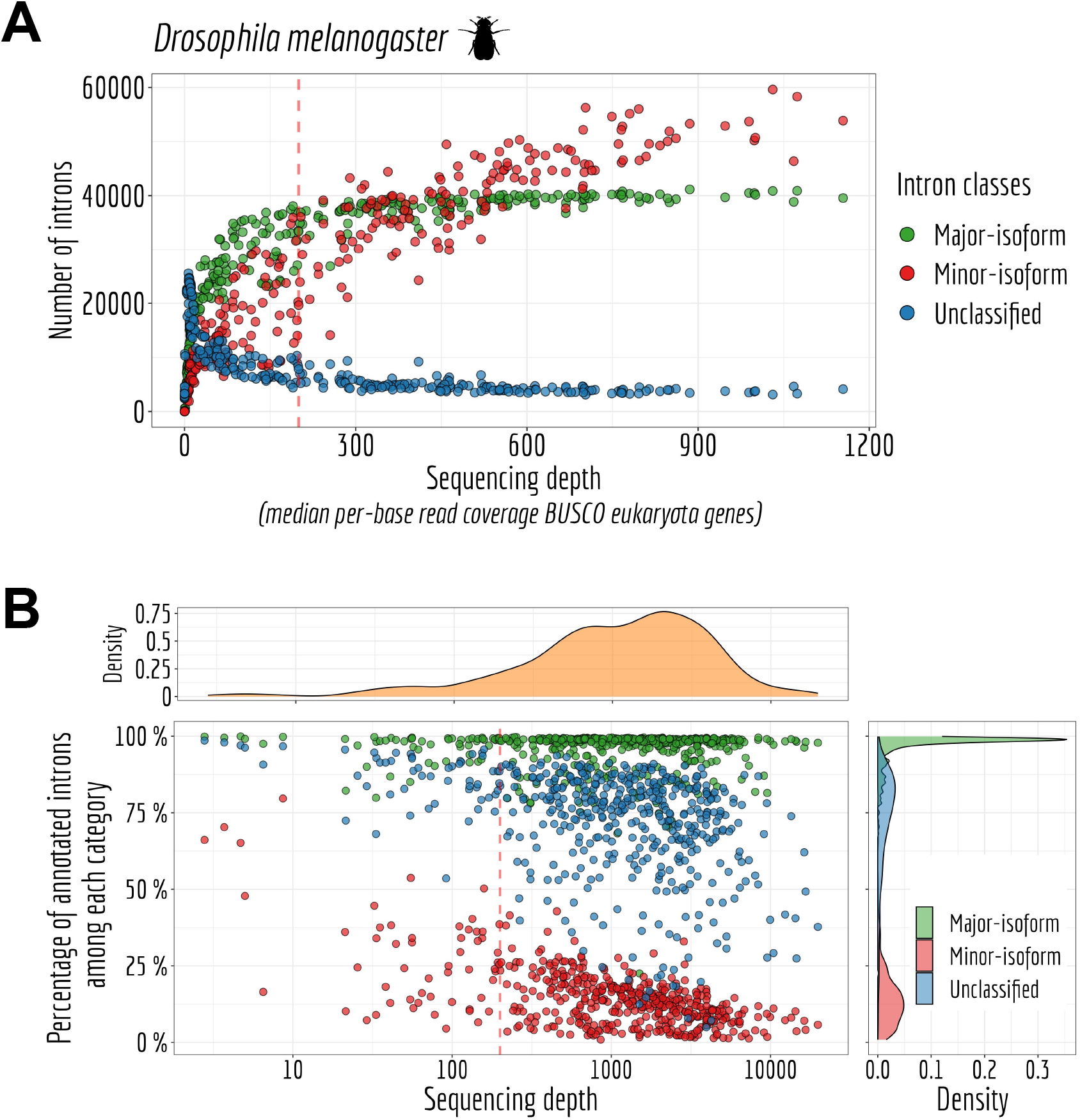
Sequencing depth impact on intron classification. **A**: Number of major (RANS *>* 0.5 and RAS *>* 0.5), minor (RANS ≥ 5% or RAS ≥ 5%) and unclassified introns for *Drosophila melanogaster*. The sequencing depth is measured by taking the median *per*-base read coverage across BUSCO genes from eukaryota gene set. **B**: *Per* species major-isoform introns, minor-isoform introns and undetermined introns (N_s_) ≥ 10) annotated proportion and sequencing depth measured by taking the median *per*-base read coverage eukaryota BUSCO genes.

As expected, we observed that the number of major-isoform introns that could be identified increased with greater sequencing depth until it reached a threshold of 200 read coverage *per* base (Fig. 9A). Beyond this threshold, no additional major-isoform introns are discernible. Simultaneously, the count of unclassified introns decreased to nearly zero, indicating that introns newly detected above the 200-read coverage threshold predominantly consisted of minor-isoform introns that shared a boundary with a major intron. Indeed, the count of minor-isoform introns continued to rise steadily beyond this point.

We then assessed the proportion of annotated introns that fall within the categories defined above. Our results reveal that the majority of species exhibit well-annotated major-isoform introns, indicating the accuracy of the intron annotation (Fig. 9B). Additionally, as sequencing depth increases, we observed a decreasing fraction of annotated minor-isoform introns. This trend is consistent with expectations, given that higher sequencing depth expands the pool of rare variants and potential spontaneous errors that may not have been previously observed. It is important to note that there appears to be no inherent limit to this phenomenon, as the intricacies of alternative splicing machinery can give rise to unpredictable errors (Bénitìere *et al*., 2024).

## Discussion

GTDrift is a comprehensive data resource facilitating investigations of genomic and transcriptomic characteristics alongside indicators of genetic drift intensity for distinct species. Notably, this resource offers information on life history traits, including longevity, adult body length, and body mass, for a curated set of 979 species. Additionally, it provides estimates of the ratio between the rate of non-synonymous substitutions over synonymous substitutions (*dN/dS*) for 1,324 species and a polymorphism-derived *N* _e_ estimates for 66 species.

For individual species, intron-centered alternative splicing frequencies, gene expression levels, and sequencing depth statistics have been systematically quantified and shared, encompassing more than 15,935 RNA-seq samples across 491 species. To enable cross-species comparisons, orthology predictions for conserved single-copy genes are provided, based on BUSCO gene sets, encompassing a total of 1,506 eukaryotic species, including 1,413 animals and 93 green plants, along with phylogenetic trees to account for phylogenetic inertia.

The number of species per data type varies due to different limitations: availability of life history traits data; completeness of gene annotations for *dN/dS* calculation; computational resources and availability of RNA-seq samples for transcriptomic analysis (Fig. 4).

These pre-processed data streamlines the work for those interested in investigating the impact of drift on biological processes across a wide range of species. All data are provided in flat files, which enable downstream computational analyses and render GTDrift mainly aimed at users with some computational skills. Nonetheless, to enhance accessibility, we have developed a user-friendly Shiny app that facilitates database exploration and allows for species-specific data downloads such as BUSCO annotation, gene expression profile, or intron splicing events (available at https://lbbe-shiny.univ-lyon1.fr/ShinyApp-GTDrift/).

### Cautionary considerations in utilizing *N* _e_ proxies

Users should bear in mind that the scientific community has yet to establish the most adequate proxies for effective population size. A prominent hypothesis suggests that these proxies are associated with the number of individuals (*N*). Indeed, species with greater longevity and larger body mass tend to be less abundant within their ecological niche due to resource (mass) and spatial (length, mass) requirements (Damuth, 1981; Nee *et al*., 1991; White *et al*., 2007). Therefore, variations in life history traits should correspond to variations in the number of individuals (*N*), which subsequently impact *N* _e_.

When using the *dN/dS* ratio as a proxy for *N* _e_, rather than focusing on correlations with the population census, we evaluated the efficiency of natural selection to purge deleterious mutations. This efficiency can be represented as the product of *N* _e_ and *s*, which denotes the selection coefficient. The extent to which a well-estimated *dN/dS* ratio can be considered as a proxy for *N* _e_ remains a subject of debate. Notably, when the rate of synonymous substitutions (*dS*) exceeds 1, it indicates a point of saturation where multiple substitutions occur *per* site, rendering *dS* susceptible to considerable noise due to the challenge of accurately identifying the number of substitutions at given sites. In such cases, the *dN* component can often still be reliably determined. Given that non-synonymous substitutions have a lower rate compared to synonymous ones, *dN* reaches a saturation point at a later stage.

Moreover, when the evolutionary time frame is relatively short, characterized by small *dS* values, the variants under examination are primarily attributed to polymorphism rather than fixed substitutions. In such cases, we are not effectively measuring substitution rates. Consequently, the discussion also revolves around determining a divergence threshold, above which we could assume that *dS* and *dN* predominantly represent substitutions, with minimal influence from polymorphism. In this perspective, the expanding polymorphism data could potentially serve as a means to distinguish between polymorphism and substitutions, offering a more efficient approach to investigate *dN/dS* (Mugal *et al*., 2014).

Overall, we found that the various *N* _e_ proxies were significantly correlated, even when accounting for the underlying phylogenetic structure. Thus, our dataset, which encompasses information on *dN* and *dS* across all branches of the phylogenetic trees, holds the potential to estimate the long-term effective population size (*N* _e_) and its interaction with life history traits over time.

### Comparing transcriptomic data

In our study, we have identified BUSCO genes for the eukaryota, metazoa, or embryophyta BUSCO reference gene sets. To ensure meaningful comparisons between species with a sufficient number of detected BUSCO genes, we evaluated the median RNA-seq coverage of these BUSCO genes. As demonstrated in Data quality validation, the median *per* -base read exonic RNA-seq coverage of BUSCO genes is a good indicator of the power to detect alternative splicing patterns. We believe that, for the inclusion of additional species, an examination of the RNA-seq read coverage on BUSCO genes is needed to ensure that we could identify major-isoform introns and analyze alternative splicing patterns.

Additionally, it is essential to assess the completeness of the genome and of the annotation, which can be estimated based on the number of identified BUSCO genes. Some species may have a limited number of well-annotated BUSCO genes, or global gene duplications may result in the presence of two copies of a BUSCO gene, which no longer qualifies as a single copy gene.

Our RNA-seq description table offers users access to information collected from the Sequence Read Archive (SRA) for the RNA-seq datasets under study. This table enables users to filter and select RNA-seq data that align with their specific research needs. Users can tailor their selection based on factors such as sex, tissue, or protocol. Depending on the research question that is asked, it may be important to extract and analyse RNA-seq samples that were generated for the same biological conditions. We provide this information so that GTDrift users are able to filter the data as needed.

To facilitate cross-species comparisons, especially in the context of alternative splicing and gene expression, users can make use of BUSCO gene sets, which should exhibit consistent expression patterns, functionality, and evolutionary constraints across diverse species. However, users should thoroughly validate this assumption and proceed with vigilance.

## Conclusion

In conclusion, we are confident that the GTDrift database can be a valuable resource for studies aiming to investigate the relationship between the intensity of genetic drift, genomic and transcriptomic characteristics.

## Acknowledgements

We thank Loïc Guille for his contribution to an initial pilot study, Tristan Lefébure for insightful discussions and Laurent Guéguen for his help on *dN/dS* analyses. Computational analyses were performed using the computing facilities of the CC LBBE/PRABI and the Core Cluster of the Institut Français de Bioinformatique (IFB) (ANR-11-INBS-0013). Thanks to Stéphane Delmotte for his wonderful help related to data storage, Shiny app deployment, calculation on cluster. Silhouette images of taxonomic Families originate from PhyloPic developed and maintained by Mike Keesey available at https://www.phylopic.org/.

## Author contributions statement

F.B. conceived the pipeline and conducted the analyses. F.B. and A.N. drafted the manuscript. All authors reviewed the manuscript.

